# Working smarter, not harder: silencing *LAZY1* in *Prunus domestica* causes outward, wandering branch orientations with commercial and ornamental applications

**DOI:** 10.1101/2024.07.03.601917

**Authors:** Andrea R Kohler, Courtney A Hollender, Doug Raines, Mark Demuth, Chris Dardick

**Affiliations:** Department of Horticulture, Michigan State University, East Lansing, MI, USA; Appalachian Fruit Research Station, Agricultural Research Service, United States Department of Agriculture, Kearneysville, WV, USA

**Keywords:** LAZY1, plum, plant architecture, gravitropism, orchard management, tree fruit production

## Abstract

Controlling branch orientation is a central challenge in tree fruit production, as it impacts factors as diverse as light interception, pesticide use, fruit quality, yield, and labor costs. In an attempt to modify branch orientation, growers use many different management practices, including tying branches to wires or applying growth regulator sprays. However, these practices are often costly and ineffective. In contrast, altering the expression of genes that control branch angles and orientations would permanently optimize tree architecture with minimal management inputs. One gene implicated in branch angle control is *LAZY1*, which promotes upward branch growth in response to gravity. We used an antisense vector to silence *LAZY1* in plum (*Prunus domestica*). We found that these *LAZY1*-silenced lines have significantly increased branch and petiole angles. In addition, they lack apical dominance and display a “wandering” or weeping branch trajectory. Given these phenotypes, we assessed whether the strength or stiffness of the branches were compromised. No differences were observed in new growth. While the wood of first-year *LAZY1*-silenced branches was more flexible and weaker than the control, the strength and stiffness of the branches were not decreased because branch diameter is increased. Finally, we evaluated the utility of *LAZY1*-silenced trees for two planar orchard systems, training them in super slender axe and espalier. *LAZY1*-silenced trees had more open canopies and were easier to constrain to the trellis height. This work illustrates the power of manipulating gene expression to optimize plant architecture for each horticultural application.

## Introduction

Controlling the orientation of lateral organs is crucial to a plant’s ability to survive and compete with other plants. In a horticultural or agricultural context, lateral organ angle impacts essential traits such as light interception (through positioning of branches), drought tolerance (through positioning of lateral roots), and harvest method (through positioning of the inflorescence). Wide branch angles are considered desirable in tree fruit production, because wide angles generally reduce bark inclusion, which leads to stronger branch unions and better ability to support fruit load^1^. Highly branched canopies with wide angles are particularly desirable in some types of high-density, commercial training systems, where the focus is on producing a narrow and uniform canopy that can support a high crop load^2^.

Due to the influence of branch angles on production, growers often expend considerable labor trying to control the position of branches through pruning, tying, and applying growth regulators^2,3^. However, as the angle of lateral organs is largely determined by genetics, these attempts can be futile. Some species, such as *Prunus domestica* (European plum), exhibit highly vigorous, upright growth habits, making them particularly difficult to train in high density planar systems^2^. In contrast to cultural practices, using selective breeding or gene editing to manipulate expression of genes that control lateral organ orientation provides a permanent solution to branch angle control throughout the lifetime of the plant^2^.

One gene family that controls lateral organ angle is the *IGT* family. The *IGT* family is named for a short, conserved (GϕL(A/T)IGT) amino acid motif ^4,5^. The family includes *TAC1* homologs, which promote outward lateral organ orientation, and *LAZY* homologs, which promote upward lateral organ orientation^6,7^. Most species have multiple paralogs of *LAZY*, which can be divided into three clades: *LAZY1*-like, *DRO1*-like, and *LAZY5*-like^4^. These paralogs are partially redundant in controlling shoot and root angles, but show distinct expression patterns with *LAZY1* homologs the most influential in shoots^8^.

*LAZY* homolog mutants across plant clades have wider lateral organ angles^8–15^. They also exhibit slowed or absent gravitropic response, sometimes to the point that shoots grow prostrate on the ground—as in *Oryza sativa* (rice)^16^, *Zea mays* (maize)^11^, and the *Arabidopsis thaliana* (arabidopsis) *lazy1,2,4* mutant^13^. In some species, *lazy* mutants also have roots that are negatively gravitropic, as in *Medicago truncatula*^12^, *Lotus japonicus*^14^, and the arabidopsis *lazy2,3,4* multiple mutant^12^. This suggests that LAZY proteins promote narrowed lateral shoot and root angles in response to gravity. Consistent with this, overexpression of some *LAZY* homologs led to narrower branch and root angles and enhanced gravitropism^17–19^. *LAZY* homologs also are involved in responses to light quantity and quality, likely functioning to integrate light and gravity signaling during the establishment of the default lateral organ angle or “set-point angle” ^15,20,21^.

The function of LAZY proteins in the gravitropic pathway has been extensively explored during the last two decades. Upon gravistimulation, LAZY proteins are phosphorylated by the kinases MKK5 and MPK3. This promotes LAZY binding to TRANSLOCON AT THE OUTER CHLOROPLAST ENVELOPE (TOC) proteins on amyloplasts^22^. As a result, LAZY proteins follow the sedimentation of the amyloplast to the new lower side of the cell, which is believed to locally enrich them in the plasma membrane ^22,23^ Once polarized to the lower side of the cell, LAZY proteins recruit the PIN3 auxin efflux carriers, likely through interactions with RCC1-like domain (RLD) proteins, which control polar auxin transport during gravitropism and development^24^. The PIN3 proteins move auxin laterally in the stems and roots,establishing a gravitropic auxin gradient with higher auxin levels on the lower side of gravistimulated roots and shoots. In gravistimulated arabidopsis *lazy1;lazy2;lazy4* triple mutants, PIN3 mis-localizes to the upper side of the cell, causing an inverted auxin gradient^25^.

Here, we present the results of a long-term field study of transgenic European plum lines with *LAZY1* silencing (*LAZY1*-sil plum) grown on their own roots and commercial rootstocks. We report control of branch angle and trajectory and pleiotropic phenotypes that may impact production. These pleiotropic phenotypes have not been previously reported, and suggest that *LAZY1* plays unexplored roles in wood development and photosynthetic pathways. Additionally, we present the results of training *LAZY1*-sil plums in planar commercial orchard systems. In this work, we not only characterize the phenotypes of these transgenic trees, but also look at how these phenotypes may impact use of these trees in production systems. This provides an example of how genetic manipulation of architecture can be used in agriculture to increase efficiency and sustainability.

## Results

### *LAZY* genes in plum

European plum is a hexaploid likely arising from an interspecific cross between *P. cerasifera* (2x) and *P. spinosa* (4x)^26^. As *P. spinosa* may itself be an allotetraploid, there may be up to three distinct subgenomes in *P. domestica*^26^. Depending on how heterozygous each subgenome is, there could be as many as six distinct homologs for each of the four *LAZY* genes previously identified in peach-- *PpeLAZY1* (Prupe.1G222800), *PpeLAZY2* (Prupe.3G308500), and *PpeDRO1* (Prupe.3G038300), *PpeDRO2*^7^. Using BlastP, we identified 20 *LAZY* homologs in plum (*PdoLAZY*), with 4-6 homologs for each *LAZY* gene (Figures 1 and S1). Of the six homologs that cluster with *PpeLAZY1*, four had an identical predicted amino acid sequence to PpeLAZY1, the fifth had a single amino acid substitution (T48N), and the sixth was either mis-sequenced, mis-annotated, or truncated, as the protein sequence started at amino acid 288 of the peach/plum LAZY1 consensus sequence (Figure S1).

**Figure 1.**
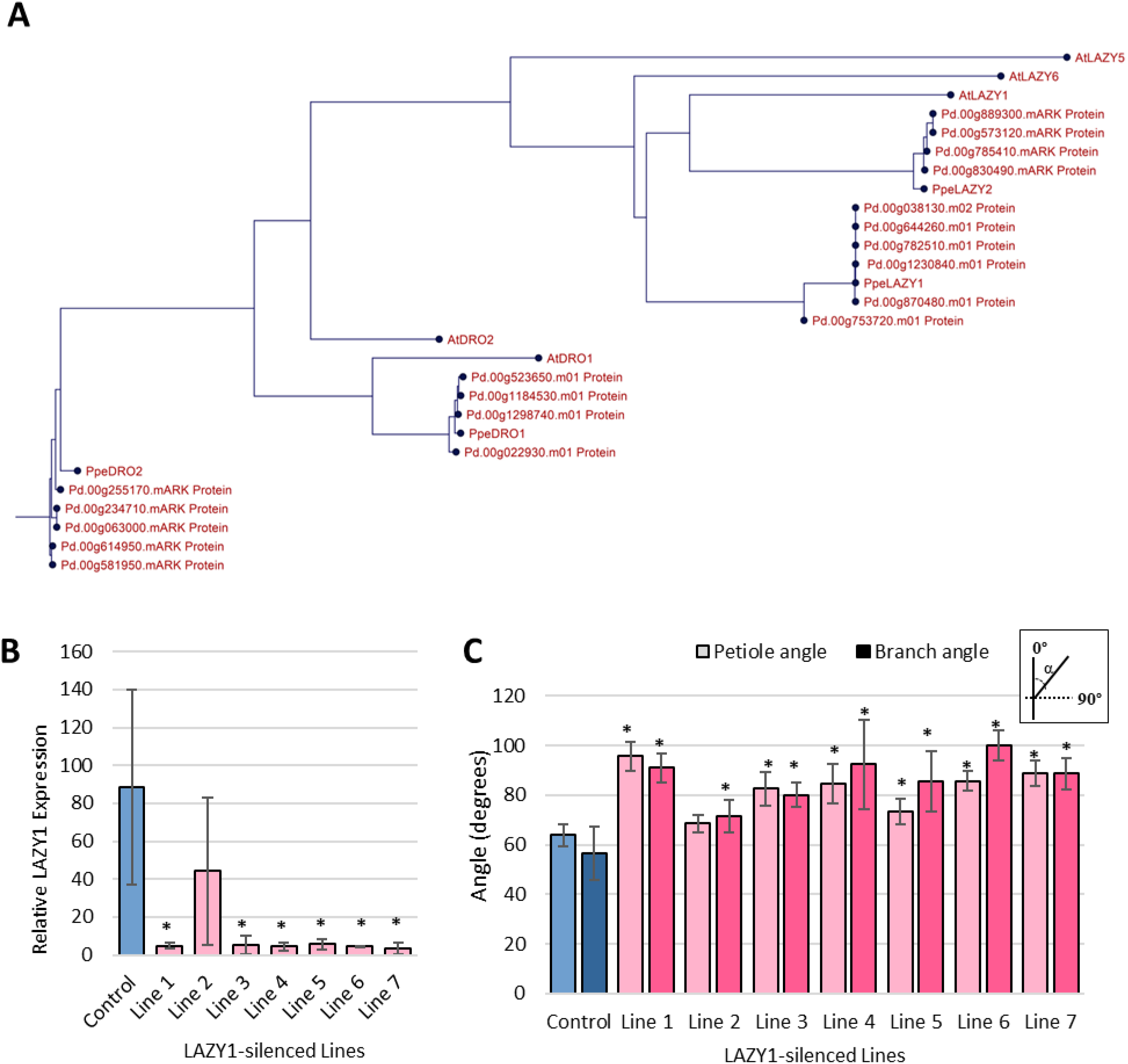
*Prunus domestica* (plum) trees transformed with a *LAZY1* antisense construct exhibited reduced expression of *LAZY1* along with wider petiole and branch angles. (A) Phylogenetic tree of *LAZY* genes in plum (Pd) used to identify LAZY1 homologs. (See Supplemental Table S1 for gene IDs and protein sequences.) Peach (*Prunus persica;* Ppe) and arabidopsis (*Arabidopsis thaliana;* At) were used for reference. (B) *LAZY1* gene expression in control and *LAZY1*-silenced lines. Expression was determined by qPCR on at least three biological replicates (trees) per line, each with three technical replicates. Values are in relation to a standard curve of known RNA from control plants. (C) Average branch and petiole angles for LAZY1-sil and control lines. Both petiole and branch angles represent the average of at least three trees per line. Diagram in the upper right indicates that angles reported are those between the branch or petiole and the apex of the shoot from which it emerged. Bars represent standard deviation and * indicates a significant difference between the control plants and a *LAZY*1-sil line (p < 0.05) according to a Student’s t-test. For B and C, control trees were plum seedlings from same cultivar that do not contain the *LAZY1-silenced* vector.

### *LAZY1*-silenced lines exhibit increased lateral organ angles and undirected lateral branch growth

To silence the *PdoLAZY1*, a construct containing a single insert of a *PpeLAZY1* sequence in the reverse orientation behind the 35S promoter in pHellsgate 8.0 (Figure S2) was transformed into plum. This served as an anti-sense vector for the five full-length *PdoLAZY1* homologs. An alignment between the *PpeLAZY1* sequence used for silencing and the *LAZY* family genes identified in plum indicated that the construct should repress only the *LAZY1* homologs, as the plum *LAZY2, DRO1*, or *DRO2* genes did not have high sequence similarity in this region (Figure S3). Seven independent plum lines were generated carrying this construct.

*PdoLAZY1* expression was quantified using qPCR primers targeting the five full-length homologs (Figure S4). *LAZY1* expression was significantly reduced in six out of seven independent *LAZY1*-sil lines (Figure 1A). These lines had significantly wider petiole and branch angles (Figures 1B and 3A). The alteration in phenotype both increased the initial crotch angle and caused a more horizontal (outward) growth trajectory (Figure 2A). Mature trees growing in both field and greenhouse environments also displayed a wandering branch phenotype, alternately arching up and down, with some branches exhibiting a weeping (downward) trajectory, and others continuing outward (Figure 2B, C). The branches appeared undirected by light or gravity and were neither oriented upward, nor consistently weeping.

**Figure 2.**
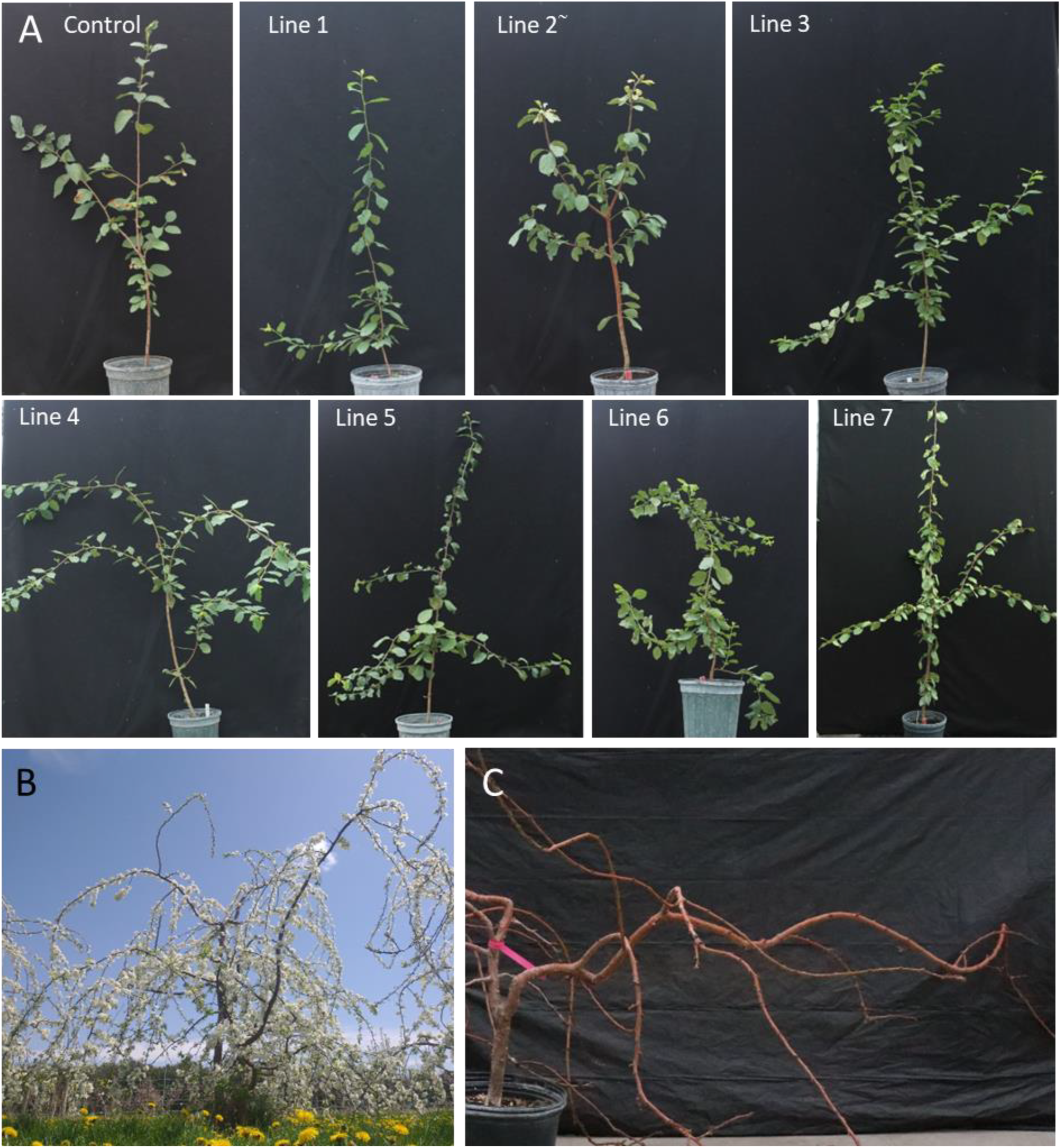
*LAZY1*-silenced plums exhibited altered leaf and branch orientations. (A) Representative control and *LAZY1-silenced* trees (1-2 years old) from each transgenic line growing in the greenhouse. ∼Note, LAZY1 expression was not significantly reduced in Line 2, as indicated in Figure 1. (B) A mature *LAZY1-silenced* Line 6 tree growing in the field. (C) A *LAZY1-silenced* Line 6 tree growing the greenhouse, demonstrating the wandering growth trajectory.

**Figure 3.**
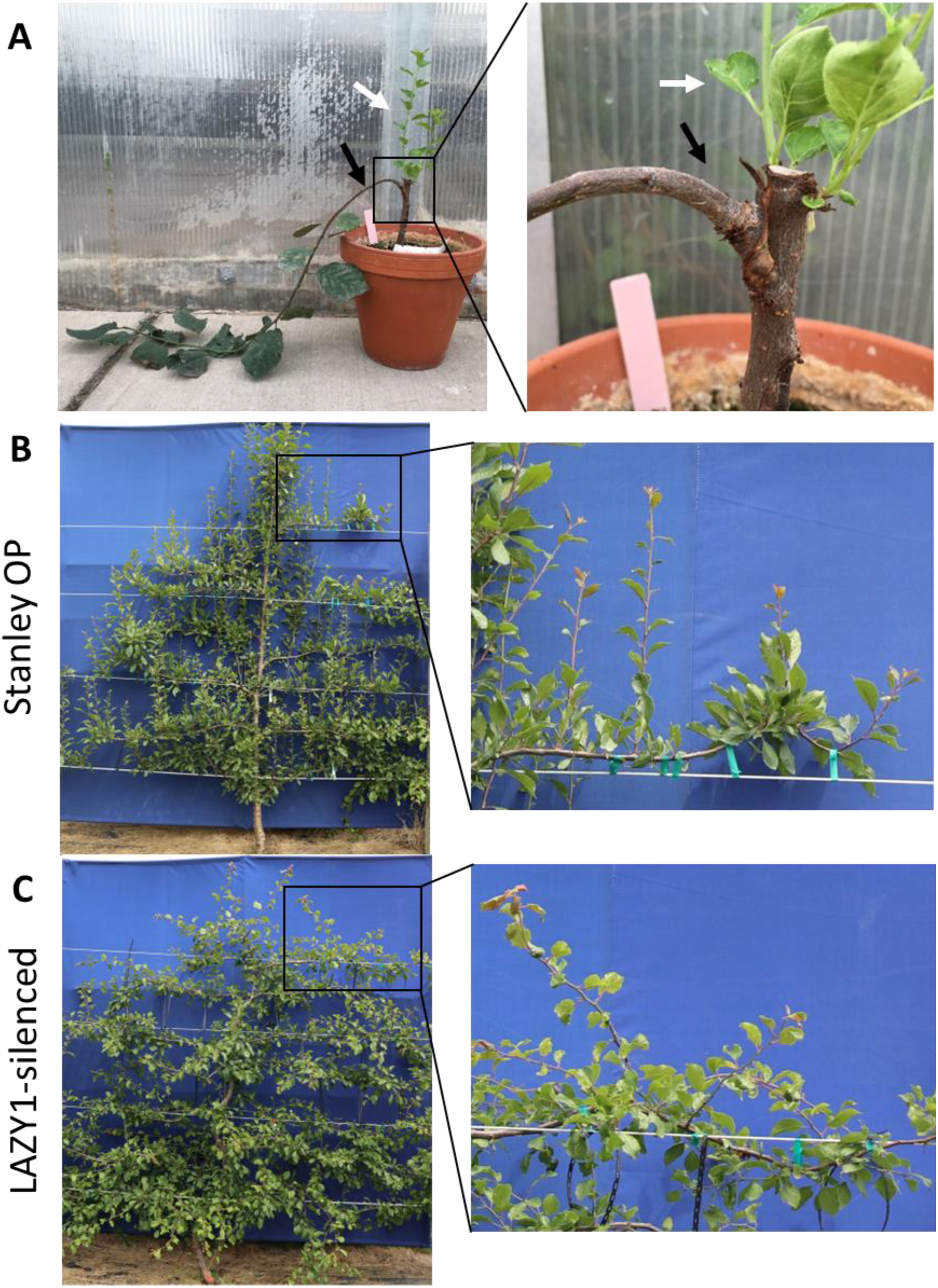
*LAZY1-*silenced shoots lacked apical dominance, even when grafted onto standard rootstock. A) Representative plum tree with a *LAZY1-*sil branch (black arrow) growing from a bud that was grafted onto a standard ‘Myrobalan’ plum rootstock, with shoots emerging from dormant rootstock buds (white arrow). B) When Stanley branches are tied horizontally, they continued trying to re-orient upwards, and buds broke from the top of the branch. C) In contrast, *LAZY1-*sil branches tied horizontal did not reorient, and buds broke from all sides of the branch.

### Horizontal shoot growth in *LAZY1*-silenced trees cannot be rescued by grafting, and trees lack gravitropic response and apical dominance

Next, we tested whether the wider emergence angle phenotypes of *LAZY1*-sil trees could be rescued by grafting. Vegetative buds from *LAZY1*-sil trees were grafted by chip-budding onto a commercial plum rootstock (‘Myrobalan’). After bud break, the shoots from the grafted buds grew horizontally, just as on the *LAZY1*-sil trees (Figure 3A). This indicated that mobile signals from the root stock could not rescue the phenotypes. Furthermore, new vegetative shoots that grew from the rootstock stem above the graft union were orientated upward. Thus, *LAZY1*-silencing in the branch below could not induce a branch phenotype through a mobile signal. (Figure 3A).

*LAZY1*-sil Line 4 and an open-pollinated ‘Stanley’ (‘Stanley’ OP) control were budded on ‘Myrobalan’ rootstock and the trees were grown in a field near Clarksville, MI. The primary shoot of each tree was tied to a stake or trellis wire to maintain vertical orientation. ‘Stanley’ OP primary shoots grew straight upward, as expected, whereas the *LAZY1*-sil primary shoots wandered and arched downward after each point where they were tied. When lateral branches were tied to horizontal trellis wire, Stanley OP branches consistently attempted to reorient growth upward (Figure 3B), whereas *LAZY1*-sil branches did not show directed growth (Figure 3C). In Stanley OP, reorienting primary branches horizontally broke the apical dominance of the tip bud, causing lateral buds to break dormancy and grow into secondary shoots. These shoots emerged primarily from the upper side of the branch and grew straight up (Figure 3B). In contrast, lateral shoots emerged from every side of the tied *LAZY1*-sil branches, and they grew in every orientation (Figure 3C).

### Branches from *LAZY1*-silenced trees do not have reduced stiffness or strength, but their wood has altered material properties

Since the LAZY1 protein is known to direct polar auxin transport through PIN3 localization, and polar auxin transport has been implicated in xylem differentiation and development^27^, we investigated the biomechanical properties of wood from *LAZY1*-sil trees by conducting materials testing experiments on new growth and one-year-old branches (Figure 4A). Neither the ability of the branch to resist bending (flexural stiffness-Figure 4B) nor the maximum force the branch can resist (F_max_-Figure 4E) was decreased in *LAZY1*-sil trees. Thus, the wandering phenotype of the branches is not due to floppiness, or an inability to hold themselves upright.

**Figure 4.**
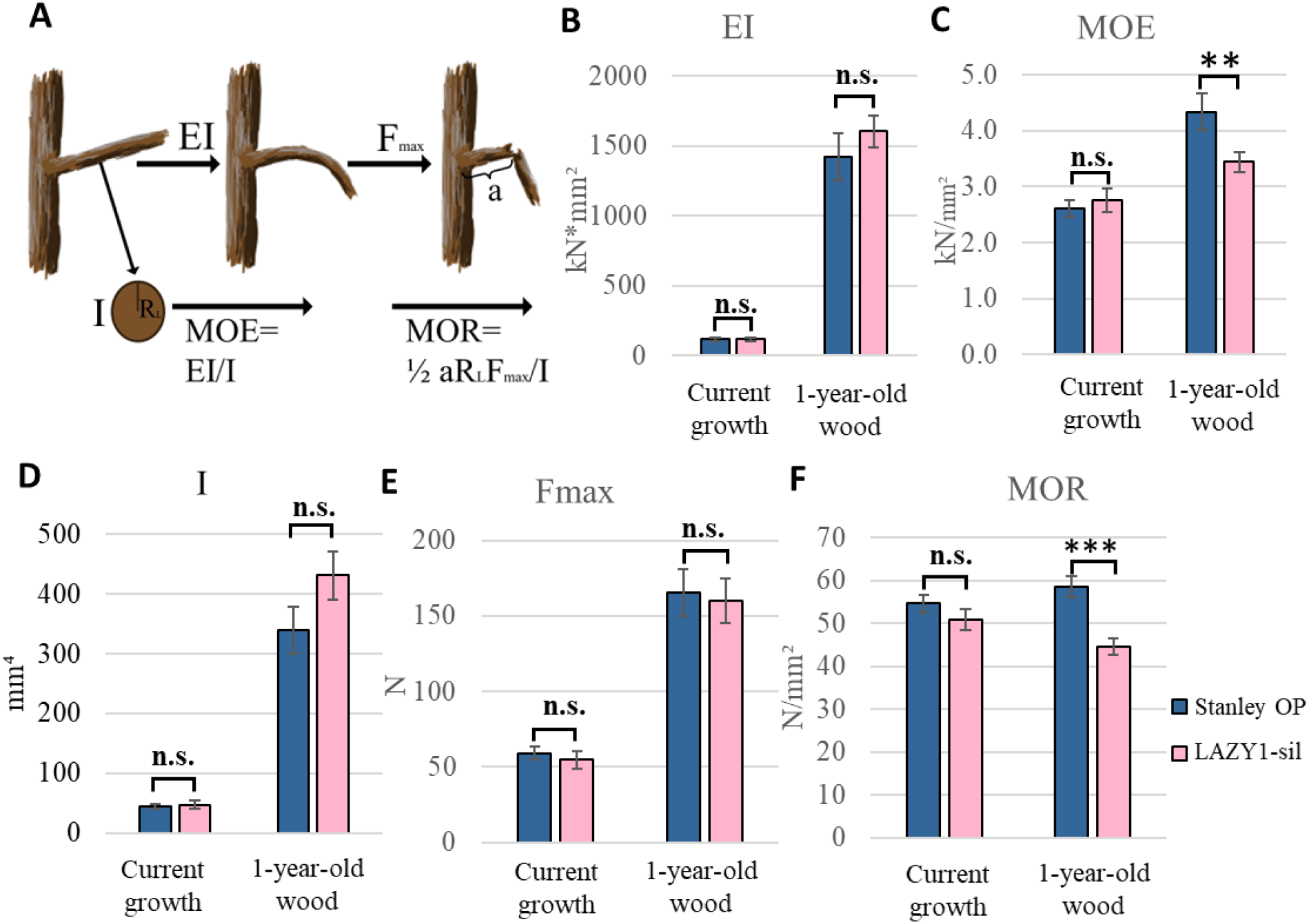
Biomechanical properties of *LAZY1-*silenced current year growth and one-year-old branches. A) Diagram illustrating biomechanical properties, including flexural stiffness (the resistance of the branch to bending, EI), area moment of inertia (a measure of how the cross-sectional area is distributed, I), radius in direction of loading (RL) modulus of elasticity (a measure of how the material resists bending in the elastic region, MOE), maximum force the branch can withstand (Fmax), distance from support to load (a), and modulus of rupture (a measure of how much force the material can withstand-MOR). Note that the formula shown for MOR is for a four-point bending test, as was performed on our branches. B-F) Biomechanical properties of *LAZY1*-silenced branches compared to ‘Stanley’: EI (B), MOE (C), diameter (D), Fmax (E), and the MOR (F) Bars represent standard error. Branches taken from 4 trees per genotype. N=30 per genotype for new growth, N=13 for ‘Stanley’ 1^st^ year, and N=14 for *LAZY1*-sil 1^st^ year. Comparisons between genotypes done with pairwise t-tests. * indicates significantly different at α=0.10, ** indicates significant at α=0.05, *** indicates significant at α=0.01, n.s. indicates not significant at α=0.10.

Despite branch stiffness and strength remaining constant, the material properties of the wood from *LAZY1-*sil branches were different than wood from ‘Stanley’ OP branches. The *LAZY1-*sil wood was more flexible (Figure 4C, lower modulus of elasticity) and weaker (Figure 4F, lower modulus of rupture). However, this did not reduce the stiffness or strength of the branches because *LAZY1-*sil branches had larger diameters, which is reflected in the increase of the area moment of inertia (Figure 4D).

### *LAZY1*-silenced leaves are chlorotic, with reduced chlorophyll and altered photosynthesis

Toward the end of each growing season, field-grown *LAZY1*-sil lines in both Kearneysville, WV and Clarksville, MI exhibited leaf chlorosis (Figures 6B-D and S5B). To quantify this, relative chlorophyll content was measured in Kearneysville for lines 1-4 in 2018 and all lines 2021 (Figures 6A and S5A). The phenotype was somewhat variable, with the extent and timing of chlorosis depending on the line, year, and location. This suggests a Genotype-by-Environment interaction in this phenotype. To investigate the potential causes and effects of the chlorosis, we measured net photosynthesis for Line 4 and Line 6, during spring and fall 2022 in the Michigan trees. Consistent with chlorophyll measurements, photosynthesis was reduced in Line 6, but not in Line 4 (Figure 5E). Interestingly, Line 6 had decreased photosynthesis at both timepoints, even though leaves were not yet chlorotic at the spring timepoint.

**Figure 5.**
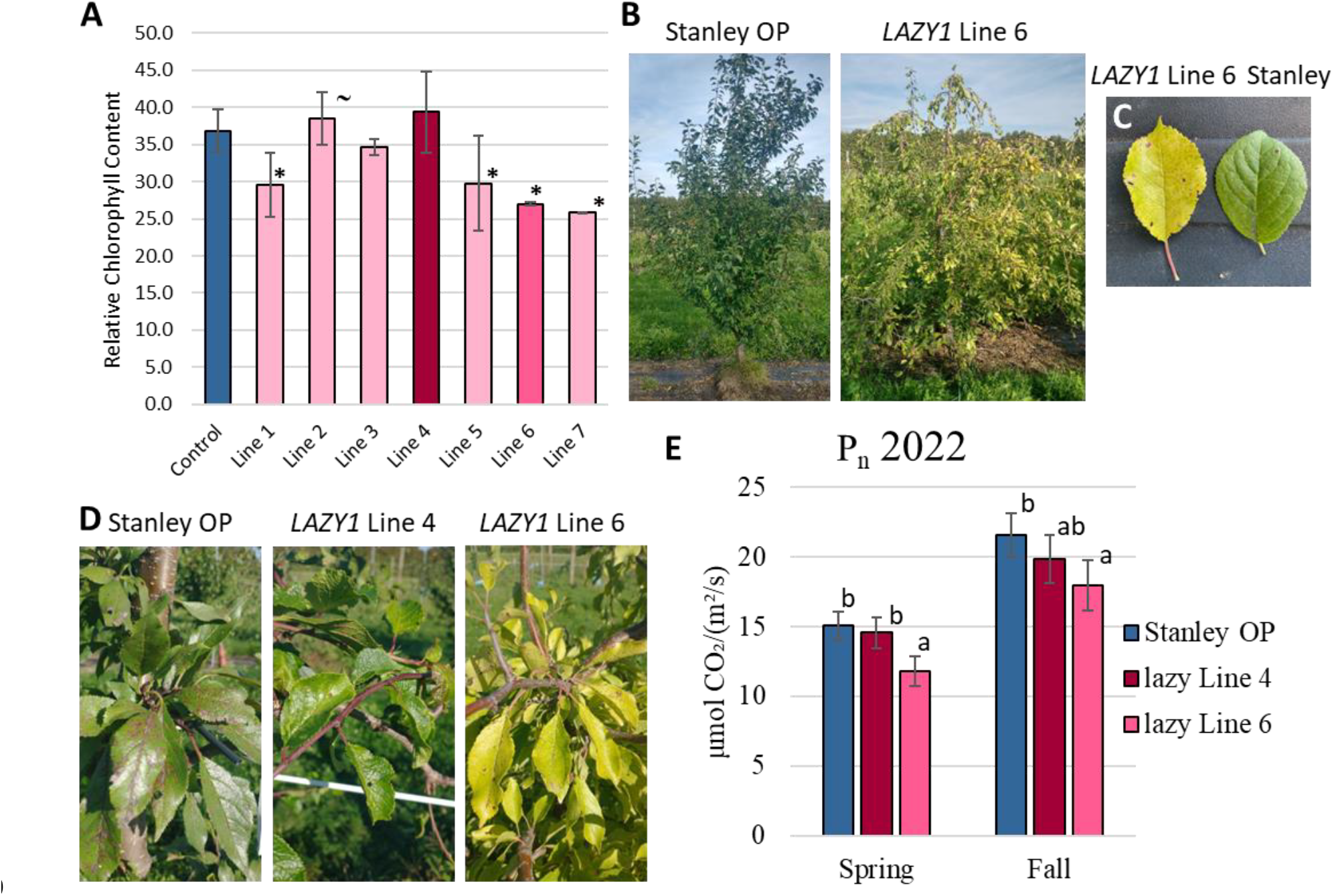
Photosynthetic phenotype of *LAZY1-*silenced. A) Chlorophyll content of *LAZY1-*sil lines in Kearneysville, WV. ∼ Line 2 did not have significant reduction in *LAZY1* expression. B) *LAZY1-*sil Line 6 and ‘Stanley’ OP control grown in Clarksville, MI. C) Leaf chlorosis comparison for ’Stanley’ and Line 6 in Kearneysville, WV. D) Leaf photos taken Fall 2021 in Clarksville, MI. E) Net photosynthesis in spring vs fall for 2022 growing season. Means within the same timepoint with the same letter are not significantly different at α=0.05. 30 or more measurements were taken for each Genotype*Timepoint combination.

**Figure 6.**
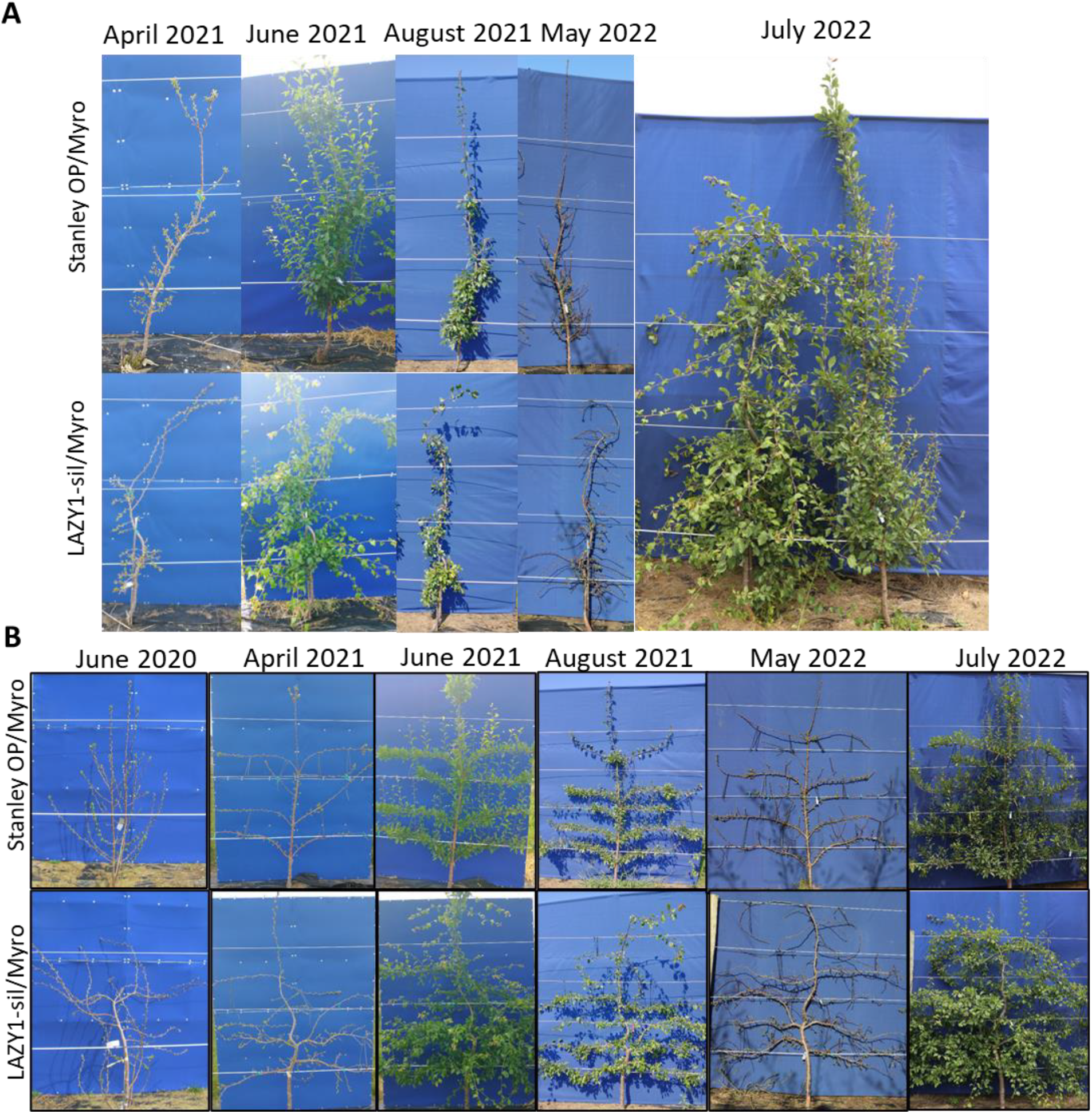
Examples of Stanley OP and *LAZY1*-silenced Line 4 on ‘Myrobalan’ rootstock trained into planar systems. A) Espalier training. B) SSA training. Note that by August 2021, ‘Stanley’ OP in both Espalier and SSA is far above the top (fourth) trellis wire, while *LAZY1-sil* is growing along it and down.

### *LAZY1*-silenced lines show unique characteristics for planar training

To test performance of *LAZY1*-sil plum trees in planar training systems, *LAZY1*-sil Line 4 trees on ‘Myrobalan’ rootstock were trained in the super slender axe (SSA) system (Figure 6A) designed for sweet cherry^28^, and the horizontal palmette espalier system (Figure 6B) used in fruit trees for hundreds of years^29^. Trees were trained and observed over the course of five years. The wider branch angles in *LAZY1*-sil lines resulted in a more open canopy, which facilitated SSA training. The wider crotch angle, horizontal orientation, and lack of gravitropic response also simplified obtaining horizontal branches in the espalier training.

‘Stanley’ trees generally sent up a single leader with limited lateral branches, which rapidly grew above the trellis (Figure 6). Heading this leader provoked a strong flush of growth at the top of the tree, shading the lower tree and draining vigor away from where it is desired lower in the tree. Because *LAZY1*-sil trunks must be staked upright for vertical growth, they do not grow significantly taller than the trellis (Figure 6). Further, because *LAZY1*-sil trees lack apical dominance, heading the main trunk does not provoke a flush of vegetative growth at the top of the tree.

## Discussion

This study demonstrated that reducing *LAZY1* expression can effectively alter branch angle in a tree fruit crop. *LAZY1-*silenced lines exhibited increased branch angle throughout the life of each plant, from the seedling stage through maturity, and the phenotype was stable when grafted onto standard rootstock. These *LAZY1-*sil lines displayed multiple traits that are beneficial for commercial production. The wider branch angles resulted in more open canopies which could increase light penetration, improving flower bud development and fruit quality^30^. An unexpected benefit of *LAZY1-* sil trees came from their lack of apical dominance. As illustrated in Figure 2B, untrained mature *LAZY1-*silenced trees had bush-like growth habits. The main trunk did not grow upright, there was no clear leader, and the trees grew more horizontally than vertically. This unexpectedly solved one of the central problems in planar systems, which is keeping the trees at or below the height of the trellis without stimulating excessive vegetative vigor by heading the tree^29^. The absence of apical dominance noticeably facilitated planar training by limiting tree height to the highest point at which it was tied, decreasing bursts of localized vegetative vigor, and increasing uniformity of the canopy. Control of vigor is particularly important in European plum, as extremely vigorous shoots generally do not produce floral buds^3^. Crucially, *LAZY1-*sil lines did not have reduced branch strength or stiffness, indicating they maintain the ability to support heavy crop loads.

These *LAZY1-*sil lines also exhibited phenotypes which may limit their applicability for commercial production but could prove useful for ornamental applications. The undirected growth of *LAZY1*-sil branches presents unique challenges for training. For Stanley espalier, horizontal branches can be trained between wires through counteracting gravitropism by tying the branches to the wire below. Since *LAZY1*-sil trees do not have gravitropic responses, it is extremely difficult to train branches between wires, as they must be tied to the wires both below and above to maintain their trajectory (Figures 4 and 8). From a training perspective, the problem can be solved by placing a wire at each interval where horizontal growth is desired and tying the branch directly to it. The wandering habit also causes idiosyncratic changes in direction of the main trunk, which makes it difficult to create a uniform canopy.

However, when allowed to develop freely, the wandering branch phenotype produces very attractive trees for ornamental use (Figure 2B). Wandering branches, as in corkscrew willows and Young’s weeping birch, are prized for adding winter interest to landscaping. Interestingly, this phenotype has not previously been reported for *LAZY1* knockdowns, even in woody species such as apple^15^. However, this phenotype may not be limited to plum. A correlative study suggested that the weeping trait in Young’s weeping birch (*Betula pendula* ‘Youngii’) is caused by a premature stop codon in a *LAZY1* homolog^31^. While this study does not describe a wandering phenotype, it is clearly visible in photographs of this cultivar available from arboretums and nurseries. This twisting phenotype is may be related to circumnutation, especially as previous studies have shown that rice *lazy1* mutants have decreased circumnutation^32^.

However, it is unclear why these windings are “frozen in time” and remain visible in the woody branches. The role of auxin transport in wood development and the changes we observed in the biomechanical properties of first-year branches from *LAZY1*-sil trees may point to a role for *LAZY1* in wood development. These changes in wood development could be what preserves and reveals the circumnutation phenotype. Alternatively, because the changes in biomechanical properties and diameter is consistent with what is observed in flexure wood or tension wood, these changes may be a secondary effect of the unique *LAZY1*-sil branch orientation, with branches forming reaction wood and/or flexure wood in response to their horizontal trajectory^33^.

Six out of seven *LAZY1*-sil lines unexpectedly showed leaf chlorosis in at least one of the seasons when chlorophyll content was measured. This chlorosis was associated with a decrease in net photosynthesis. While undesirable for production systems, their unique color may be a desirable trait for ornamental use. The variance of the phenotype among lines and seasons suggests that it didn’t have 100% penetrance, which may assist in selecting the desired phenotype. Leaf chlorosis has not been reported for *lazy1* mutants. This may be because most studies on *lazy1* mutants were short-term experiments using annual species grown in growth chambers. The chlorotic leaf phenotype in plum became apparent midway through the growing season in the high light intensity of the greenhouse and field. Given recent revelations about the ability of LAZY1 protein to associate with the TOC1 complex found in amyloplasts and other plastid membranes, as well as its role in light response/signaling^22,23^, the leaf chlorosis may indicate a role for LAZY1 in the response of photosynthesis to light-related signaling, the absence of which results in damage. This hypothesis is strengthened by the observation that photosynthesis is reduced in the spring in chlorotic Line 6, prior to any observable leaf chlorosis.

In the ongoing quest to produce tree fruit more efficiently, advances that improve tree structure and fruit yield while decreasing labor inputs will be crucial to success. Due to the substantial amounts of labor required to physically constrain and control tree canopies over the lifespan of the orchard, modifying the genetics controlling canopy structure will be a key part of long-term solutions. Here we have demonstrated that manipulation of *LAZY1* expression can permanently alter canopy structure, providing traits beneficial to planar training systems. While some of the pleiotropic phenotypes we observed were deleterious to commercial production, this work provides a strong foundation for further investigation of genetic manipulation of branch angle in tree fruit.

## Materials and methods

### *LAZY* gene identification and phylogeny

*LAZY* genes in *P. domestica* were identified using BLASTp of the previously identified peach LAZY proteins^7^ against predicted proteins from the *P. domestica* draft genome v1.0^34^(Figure 1 and Supplemental Table 1) Due to a highly unusual exon-intron structure, *LAZY* genes are often mis-annotated^4^. This structure is conserved across species, with each gene containing five exons, with the first exon including just two codons (often an “ATGAAG”) and the last exon containing only ∼20 bases^4^. To identify and correct errors in annotation, the DNA sequences of plum genes identified as *LAZY* homologs were aligned to the homologous peach and arabidopsis sequences. Using that alignment, the plum and peach genes were re-annotated with particular attention to identifying a gene model that fit the conserved gene structure. Following re-annotation, the protein sequences were predicted, and the resulting protein sequences were used for subsequent alignments.

Alignments and phylogenetic trees were produced in CLC Genomics Workbench v22.0. Alignments were performed using a gap open cost of 10.0 and a gap extension cost of 1.0, the alignment set to “Very accurate.” The protein phylogenetic tree was constructed using the Neighbor Joining method with Jukes-Cantor as the protein distance measure, and 100 bootstrap replicates.

### Cloning

To generate *LAZY1* silenced plum lines, a 306bp fragment corresponding to the peach LAZY1 gene (peach genome version 1.0 ID ppa007017, now named Prupe.1G222800 in genome version 2) was amplified using primers PpLazy-1 1F (5’ AAG CCA AAC TGT GGC ACA AAG C) and PpLazy-1 2R (5’AGC TGC CAG GAC TTT CTC CAA T), cloned into the pENTR-D TOPO vector (Invitrogen, Carlsbad, CA), and then into the pHellsgate 8.0 vector^35^ using LR Clonase (Invitrogen, Carlsbad, CA). While the intended construct would have had two copies of the 306bp fragment as a hairpin, the construct used for transformation was later discovered to only have a single insert in the reverse orientation behind the 35S promoter within pHellsgate 8.0, creating an anti-sense vector rather than an RNAi construct.

### Plum transformation

The pHELLSGATE 8.0 plasmid containing the peach *LAZY1* gene fragment was transformed into Agrobacterium tumefaciens strain GV3101. The gene construct was engineered into European plum (*Prunus domestica* L) following the protocol of Petri et al.^36^. Cold (4°C) stored seeds of ‘Stanley’ plum were used for transformation. Briefly, the seeds were first cracked to remove the stony endocarp, surface sterilized with 15% commercial bleach for 15 min., washed three times with sterile water, and the hypocotyl slices were excised from the zygotic embryos under a laminar flow hood using a stereomicroscope. After incubating for 20 min. in an *Agrobacterium* suspension, the transformed hypocotyl sections were cultured for 3 days in co-cultivation medium. Finally, the hypocotyl sections were plated in antibiotic (80 mg/l kanamycin) selection medium to select transgenic shoots. The kanamycin resistant transgenic shoots were multiplied in plum shoot multiplication medium, rooted, acclimatized in the growth chamber and planted in 15-23cm pots in a temperature-controlled greenhouse to evaluate growth and development.

### Plant material

Greenhouse-grown *LAZY1*-sil and control lines were used for quantifying gene expression and for branch and petiole angle analysis. Trees were 1-2 years old when gene expression was quantified and petiole angles were measured, and 2-3 years when branch angles were measured.

The controls for lazy expression were transformed plum seedlings from open-pollinated plum cultivar ‘Stanley’ (Stanley OP) that did not contain the *LAZY1*-sil vector. These plums instead contained the pSUC/PSUL vector, which does not impact branch angle^37^.

*LAZY1*-sil plum trees and Stanley OP controls were also planted at the USDA ARS Appalachian Fruit Research Station (AFRS) research field near Kearneysville, WV and at the Michigan State University Clarksville Research Center (CRC) near Clarksville, MI. The trees at AFRS were planted at ∼2.4 m spacing in October 2014. The *LAZY1*-sil plums were planted in two blocks of three trees. Control plums were planted between groupings of *LAZY1*-sil trees. The trees at CRC were generated in 2017 and 2018 from budwood from AFRS and were planted in September 2018. *LAZY1*-sil Line 4 trees and Stanley OP trees, grafted onto ‘Myrobalan’ rootstock, were used for training studies. The trees were planted at ∼2.4m spacings for horizontal palmette espalier training and ∼0.9m spacing for SSA training. The trees were trained to a trellis with ∼46 cm spacing between wires. The CRC planting also contains the *LAZY1*-sil Line 6 trees on their own roots.

### RNA extraction and gene expression analysis

Total RNA was extracted from frozen tissue samples using E.Z.N.A SQ Total RNA Kit (Omega Bio-tek, Inc., USA), according to the manufacturer’s instructions. Leaf tissue was used for extraction and for expression analysis in transgenic plums. The resulting RNA samples were then treated with DNase I to remove contaminating genomic DNA. To determine gene expression levels, qPCR reactions were carried out using the gene specific primers PpLAZY-qPCR-5F (5’ ATGCTTTATGCTTCTTCTCG) and PpLAZY-qPCR-5R (5’ TTGCTCAGCAGATGAGGT; Figure S3), and the SuperScript III Platinum SYBR Green One-Step qRT-PCR Kit with ROX (Invitrogen Corp., USA). The qPCR was run using an ABI 7900DNA Sequence detector (Applied Biosystems) according to the following parameters: cDNA synthesis step at 50 °C for 5 min, followed by PCR reactions at 95 °C for 5 min and 40 cycles of 95 °C for 15s, 60 °C for 30s, and a final cycle of 40 °C for 1 min. The qPCR was performed on RNA from three to four independent biological replicates (trees) for each transgenic line, as well as on five control trees. Each biological replicate had three technical replicates for the reactions. Relative expression values were determined using a standard curve (generated from serial dilutions of RNA from a tree that did not have the *LAZY1* vector), which was run at the same time.

### Petiole and branch angle measurements

Petiole angles for up to ten leaves growing from the trunk of young (<1-year old) trees that had not yet initiated lateral shoots were measured in 2013 using a protractor. For the control and lines 3 through 7, five trees were measured. Ten trees were measured for Line 1, and three trees for Line 2. Crotch (branch) angles for six branches from three to ten trees per genotype growing at AFRS were measured in 2021 using a protractor. Angles represent the angle between the branch and the shoot from which it emerged. When the branch angles were measured, the trees were approximately nine years old and had been growing in the field for six years.

For both petiole and branch angle statistical analyses, the average angle per tree was considered a biological replicate, and standard deviations are based on those averages. Significant differences between the *LAZY1*-sil lines and control trees were determined by a p-value < 0.05 from Student’s t-test.

### Biomechanics measurements

Clonal trees were generated from budwood from a single open-pollinated ‘Stanley’ (Stanley OP) tree and a single *LAZY1*-sil Line 4 tree grafted onto ‘Myrobalan’ rootstock, grown at CRC and trained as horizontal palmette espalier. Horizontally oriented new growth and 1^st^ year wood (branches initiated the previous season) were collected July 9, 2023. Four to five branches per tree were collected from four trees per genotype for new growth, for a total of 20 ‘Stanley’ OP and 19 *LAZY1*-sil samples. Three to five branches per tree were collected from three trees per genotype for 1^st^ year samples, for a total of 13 ‘Stanley’ OP and 14 *LAZY1*-sil samples.

Four-point bending tests were conducted on an Instron Universal Testing Machine (Model 4202, Instron Corporation, Canton, MA) using a 500N load cell. The span of the outer supports was set to 90mm, while the two posts of the actuator were set to 30mm. Leaves and lateral branches were removed from a section of wood ∼10 cm long, and the section was oriented so that the upper side of the branch was positioned down on the Instron, so that force was exerted in the same direction as gravitropic force acted on the branch in its original position. The force versus displacement curve was measured until a maximum force was reached.

Flexural stiffness (EI) was calculated using the equation EI=(F/V)(a^2^/12)(3L-4a), where (F/V) is the slope of the linear section of the force/displacement curve, a is the post to load distance and L is the total length of the span. Flexural stiffness was used to calculate the modulus of elasticity (MOE), using the equation MOE=EI/I where I is the area moment of inertia. For a branch with an elliptical cross section, I= (π/4) (R_P_)(R_L_^3^), where R_L_ is the radius in the direction of loading and R_P_ is the radius perpendicular to loading. The modulus of rupture (MOR) was calculated using the equation MOR=(1/2 Fmax)(a)(R_L_)/I, where Fmax is the maximum force withstood by the branch. To test for statistically significant differences between the genotypes, a Student’s t-test was performed using the T.TEST function in Excel, assuming a homoscedastic two-tailed distribution.

### Chlorophyll and photosynthesis measurements

Chlorophyll measurements were taken in triplicate from field-grown trees at the AFRS using a SPAD 502 Chlorophyll Meter (Konica Minolta, Inc., Tokyo, Japan).

Photosynthetic parameters were measured in late spring (June 9^th^ and June 23^rd^, 2022) and early fall (October 5, 2022) using a CI-340 infrared gas analyzer (CID Bio-science, Camas, WA). Measurements were taken with an open system in differential mode, with 6.25 cm^2^ of leaf area in the chamber, and a flow rate of 0.3 l/min. Measurements were taken from trees at CRC, including control trees (‘Stanley’ OP on ‘Myrobalan’ rootstock and on own root), *LAZY1*-sil Line 4 on ‘Myrobalan’, and *LAZY1*-sil Line 6 trees on their own roots. At each time point, three to five measurements were taken per leaf, on three to five leaves per tree, with two to six trees per genotype.

Statistical analysis was performed in R v4.3.1. To control for the environmental variability of the field conditions, net photosynthesis (Pn) for all genotypes from two seasons (2021 and 2022) was modeled as a response to photosynthetically active radiation (PAR), carbon dioxide concentration at intake (CO2in), air temperature (Tair), air pressure (Pressure), and water vapor pressure (H2Oin) at intake. Externally studentized residuals were used to identify influential outliers, and observations with a studentized residual greater than |2| were dropped (10 in total). Studentized residuals and the criterion Cooks Distance for point>3(mean Cooks distance) was used to identify outliers. Outliers were then manually checked. Outliers for which there were two or more outliers per leaf were retained as reflecting true biological variability, while single outliers were omitted. The model was then re-checked without outliers. Variance inflation factor (VIF) was used to check for collinearity of the environmental variables. Added variable plots were used to check for linearity and the contribution of each additional variable. Variables with the least contribution or the highest VIF were sequentially dropped until VIF<3 for all variables, and all variables had a clear contribution to the model. Finally, variables for genotype and for accounting for the repeated measures were added to the model and each year was modeled separately.

The resultant model for photosynthesis was Pn = μ + Genotype + Timepoint + Genotype*Timepoint + (1|Genotype:Tree) +(1|Genotype:Tree:Leaf) + PAR+ CO2 + Tair+ Pressure+error where μ indicates the grand mean, “Genotype” is a variable for the control or RNAi line, “Timepoint” accounts for the effects spring versus fall, “Genotype*Timepoint” accounts for the interaction between genotype and timepoint, (1|Genotype:Tree) controls for the effects of tree, which is a random variable nested within genotype, (1|Genotype:Tree:Leaf) controls for leaf to leaf variation, and is nested in tree, and “PAR”, “CO2in” and “Tair” are continuous variables controlling for their respective environmental factor. This model was gated with ANOVA, and normality and equal variances of the residuals were checked. Since interaction between genotype and timepoint was significant, all pairwise comparisons were performed for genotype slicing by timepoint using t-tests at α=0.05.

## Acknowledgments

Funding for this work was provided by Michigan State University, The Michigan State Horticultural Society, The United States Department of Agriculture National Institute of Food and Agriculture HATCH project 1013242 (CAH), MSU AgBioResearch Project GREEEN grant GR18-031, and the Agriculture and Food Research Initiative Competitive Grants Program, grant no. WVAW-2011-04220, from the USDA National Institute of Food and Agriculture (CD).

We would like to thank Srinivasan Chinnithambi and Ralph Scorza for performing plant transformations, Elizabeth Lutton for cloning the constructs, Frank Telewski for access to and assistance with his Instron, Greg Lang for help implementing planar training systems, Peter Kohler for assistance with data formatting and statistical analysis, Joy Johnson and Andrew Scheil for assistance with photosynthesis measurements and imaging, and the MSU Clarksville Research Center Farm Manager Dan Platte and his team for helping with tree planting and maintenance.

## Author contributions

CAH and CD conceived project idea. CAH, ARK, and CD planned experiments. CAH, DR, MR, and ARK performed experiments and analyzed data. ARK wrote the manuscript with input from other authors. CAH and CD edited the manuscript.

## Conflict of interest

Authors declare no conflict of interest.

## Data availability

The data underlying this article will be shared on reasonable request to the corresponding authors.

## Supplement

**Figure S1: LAZY protein alignment**. Pd=*Prunus domestica;* Ppe=*Prunus persica*; At= *Arabidopsis thaliana*.

**Figure S2: PpeLAZY1 (Prupe.1G222800) mRNA sequence**. Cloning primers are highlighted in yellow and the remaining insert sequence in gray.

**Figure S3: Plum *LAZY* gene alignment to the insert sequence from *PpeLAZY1***. Cloning primers are highlighted in yellow and the remaining insert sequence in gray.

**Figure S4: Sequence alignment for sequence used for gene expression primers**.

**Figure S5: Additional chlorosis data**. A) 2018 chlorophyll data for WV trees on 8/16/18. ∼Line 2 did not have significant reduction in *LAZY1* expression. B) Leaf phenotypes for Stanley OP control and *LAZY1* Line 4 on 7/27/23

**Table S1. Peach, plum, and arabidopsis LAZY homologs ID numbers and protein sequences**

## Notes

### Competing Interest Statement

The authors have declared no competing interest.

