## Supplemental Figures for "Working smarter, not harder: silencing *LAZY1* in *Prunus domestica* causes outward, wandering branch orientations with commercial and ornamental applications"

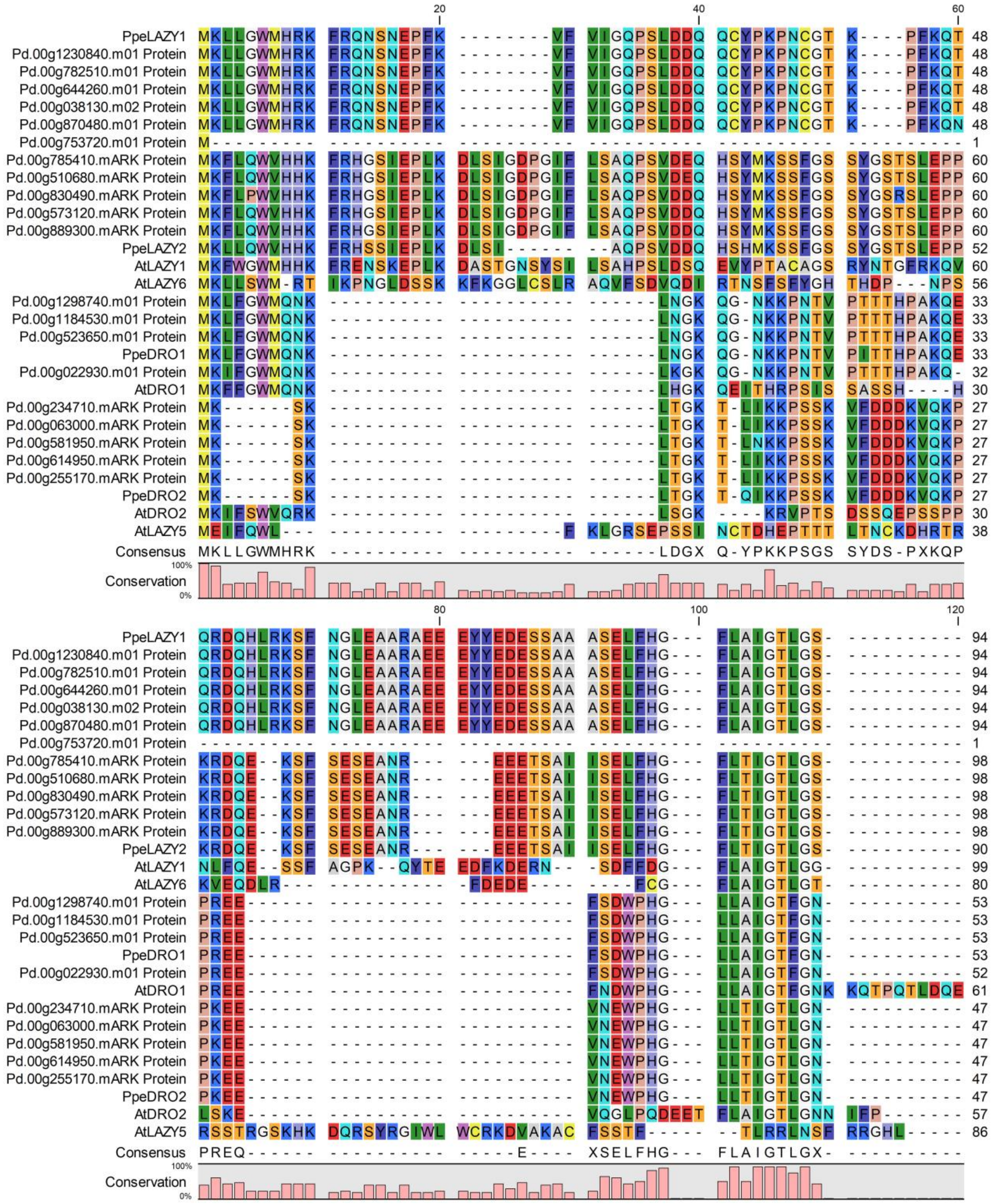

**Figure S1: LAZY protein alignment.** Pd=*Prunus domestica;* Ppe=*Prunus persica*; At= *Arabidopsis thaliana.*

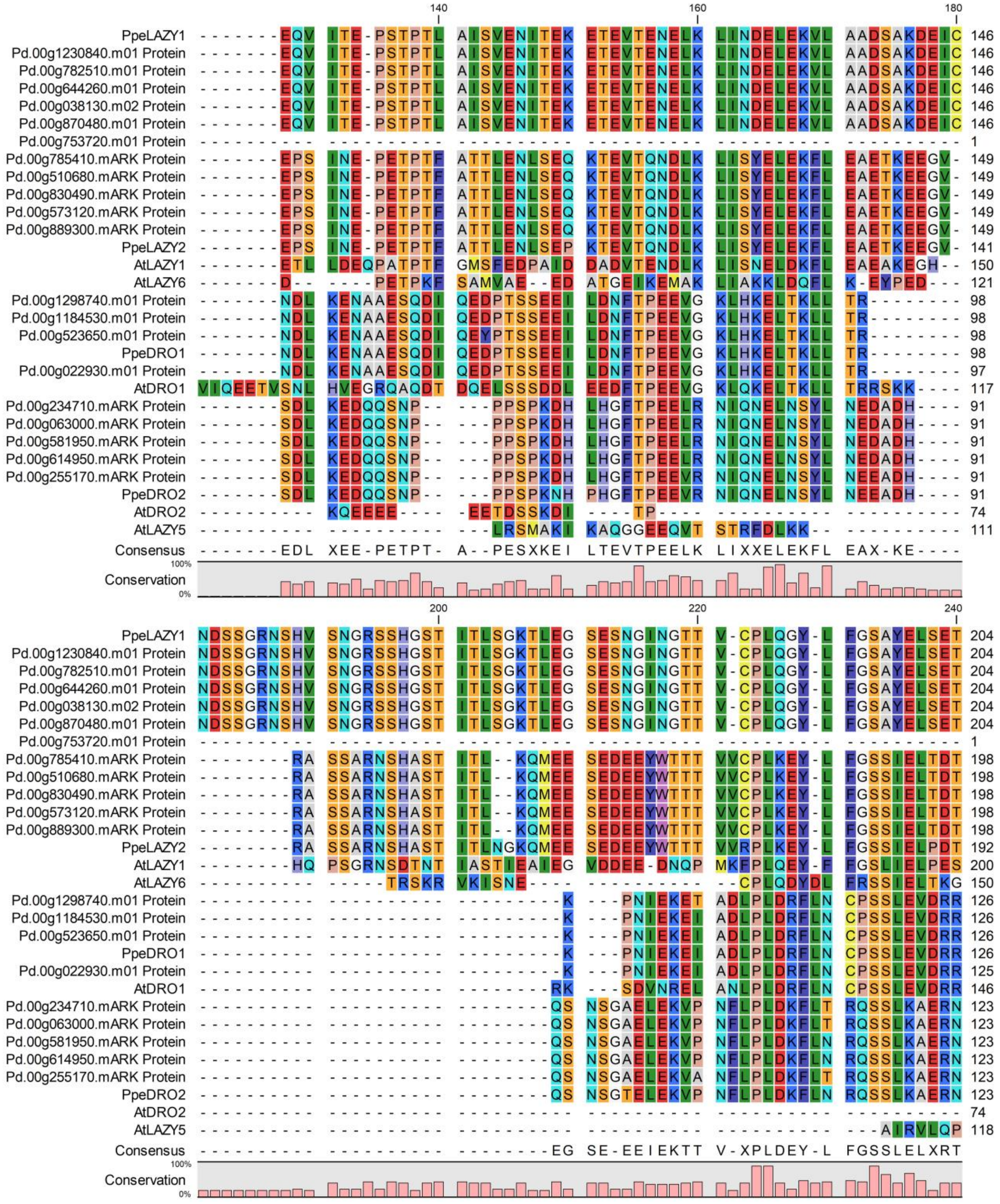

**Figure S1: cont.**

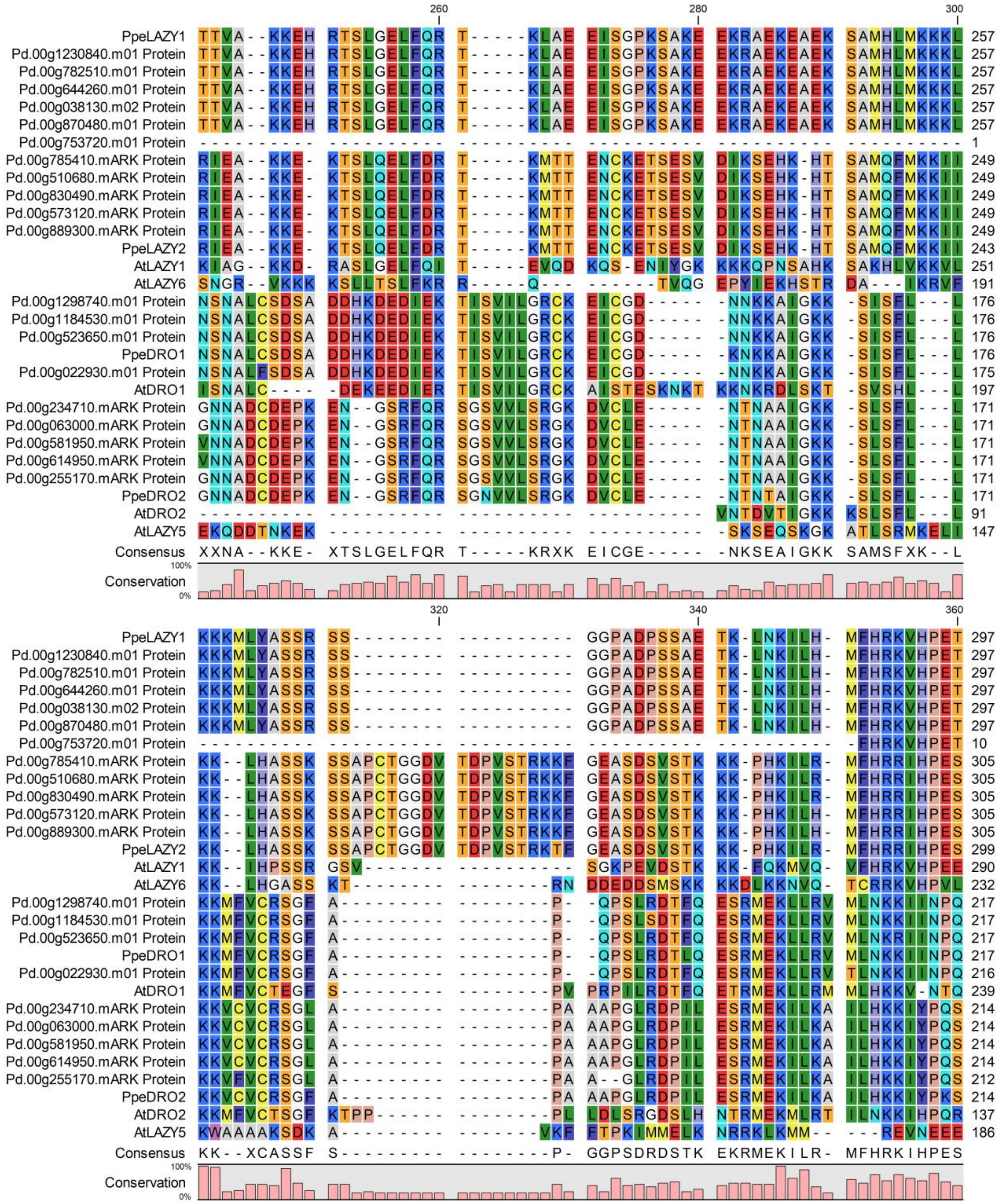


**Figure S1: cont.**

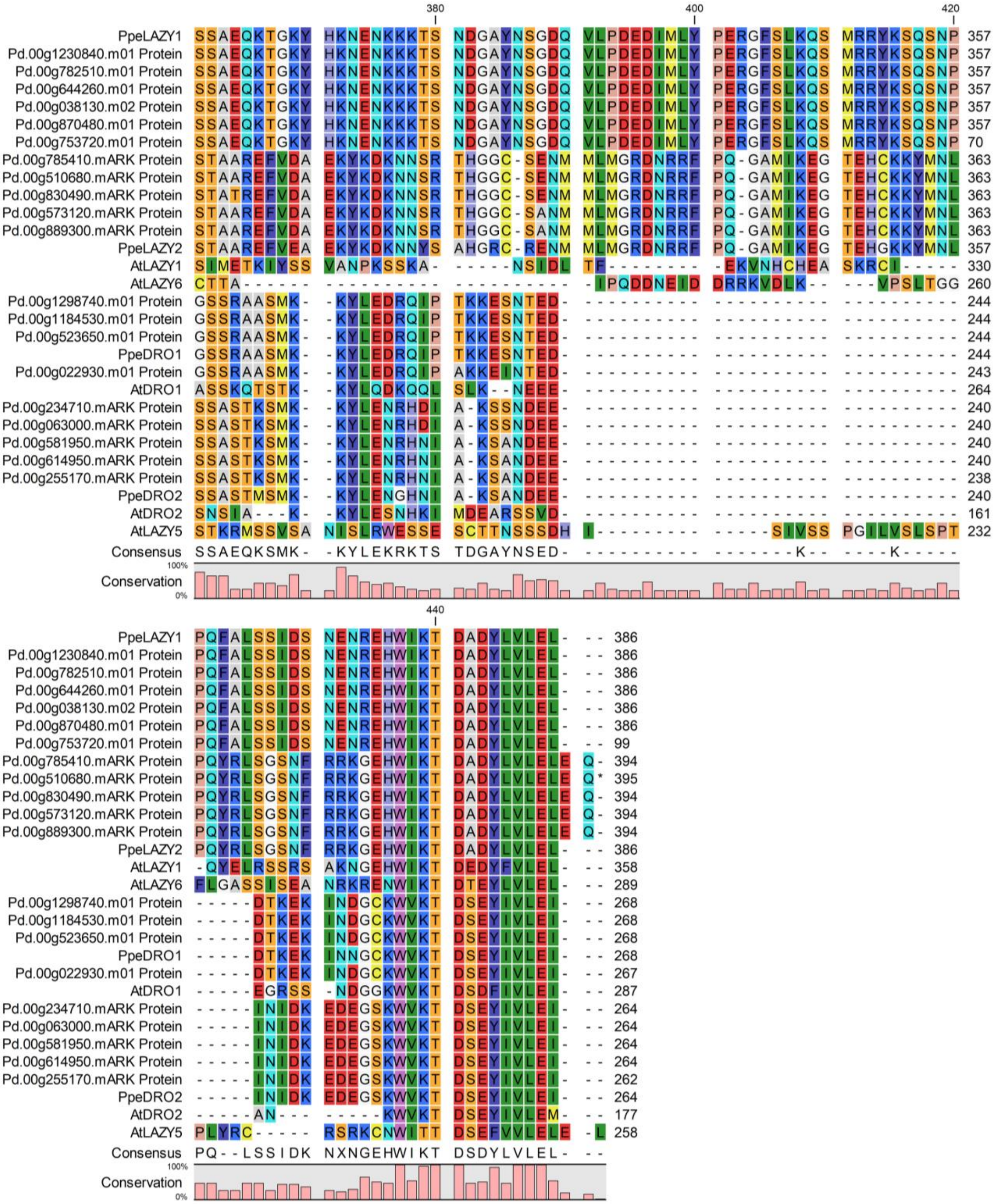


**Figure S1: cont.**

ATGCCAATGTTGCAGTTACTAGGTTGGATGCATCGTAAGTTTCGGCAGAATAGCAACGAGCCAT
TTAAAGTTTTTGTCATTGGGCAGCCATCTCTCGATGATCAACAATGCTATCCTAAGCCAAACTGTG
GCACAAAGCCCTTTAAACAAACCCAGAGAGACCAGCACCTTCGGAAGTCTTTCAACGGTCTAG
AGGCAGCCAGGGCAGAAGAAGAATACTATGAAGATGAATCATCTGCTGCAGCATCTGAGCTCT
TCCATGGCTTCCTTGCAATTGGTACCCTTGGCTCAGAGCAAGTCATCACGGAACCATCAACTCCA
ACACTTGCCATCTCTGTGGAGAACATAACTGAAAAAGAGACTGAGGTCACAGAGAATGAATTG
AAGCTCATCAATGATGAATTGGAGAAAGTCCTGGCAGCTGATTCAGCTAAAGATGAGATTTGCA
ATGATTCATCTGGAAGAAACAGCCATGTTAGCAATGGAAGAAGTAGCCATGGTAGCACCATCAC
ACTAAGTGGCAAGACACTGGAAGGCTCAGAGAGCAATGGGATTAATGGAACCACAGTGTGCC
CACTCCAGGGATATCTTTTTGGGTCAGCATATGAATTGTCAGAAACAACAACAGTGGCAAAGAA
GGAACACAGGACATCTCTTGGCGAGCTGTTTCAGAGGACTAAATTGGCAGAGGAGATTTCTGG
ACCGAAATCGGCCAAGGAGGAGAAGCGAGCAGAGAAGGAAGCTGAAAAGTCCGCCATGCAC
TTGATGAAAAAGAAGCTCAAGAAAAAAATGCTTTATGCTTCTTCTCGCAGCTCTGGTGGACCTG
CAGATCCTTCCTCAGCGGAAACAAAACTGAATAAGATCCTTCACATGTTCCACAGAAAAGTTCA
CCCTGAAACCTCATCTGCTGAGCAAAAAACTGGTAAGTACCATAAGAACGAAAACAAGAAGAA
AACAAGCAATGATGGGGCTTACAACAGTGGAGATCAGGTGCTTCCAGATGAAGACATCATGCT
ATATCCTGAACGAGGCTTCTCCTTGAAGCAGAGCATGCGGCGCTACAAGAGTCAATCAAACCCA
CCGCAATTCGCGCTTAGCAGCATTGATTCAAATGAGAACAGGGAGCACTGGATCAAAACAGAT
GCAGACTACTTAGTCTTGGAGCTGTGA

**Figure S2: PpeLAZY1 (Prupe.1G222800) mRNA sequence.** Cloning primers are highlighted in yellow and the remaining insert sequence in gray. **.**

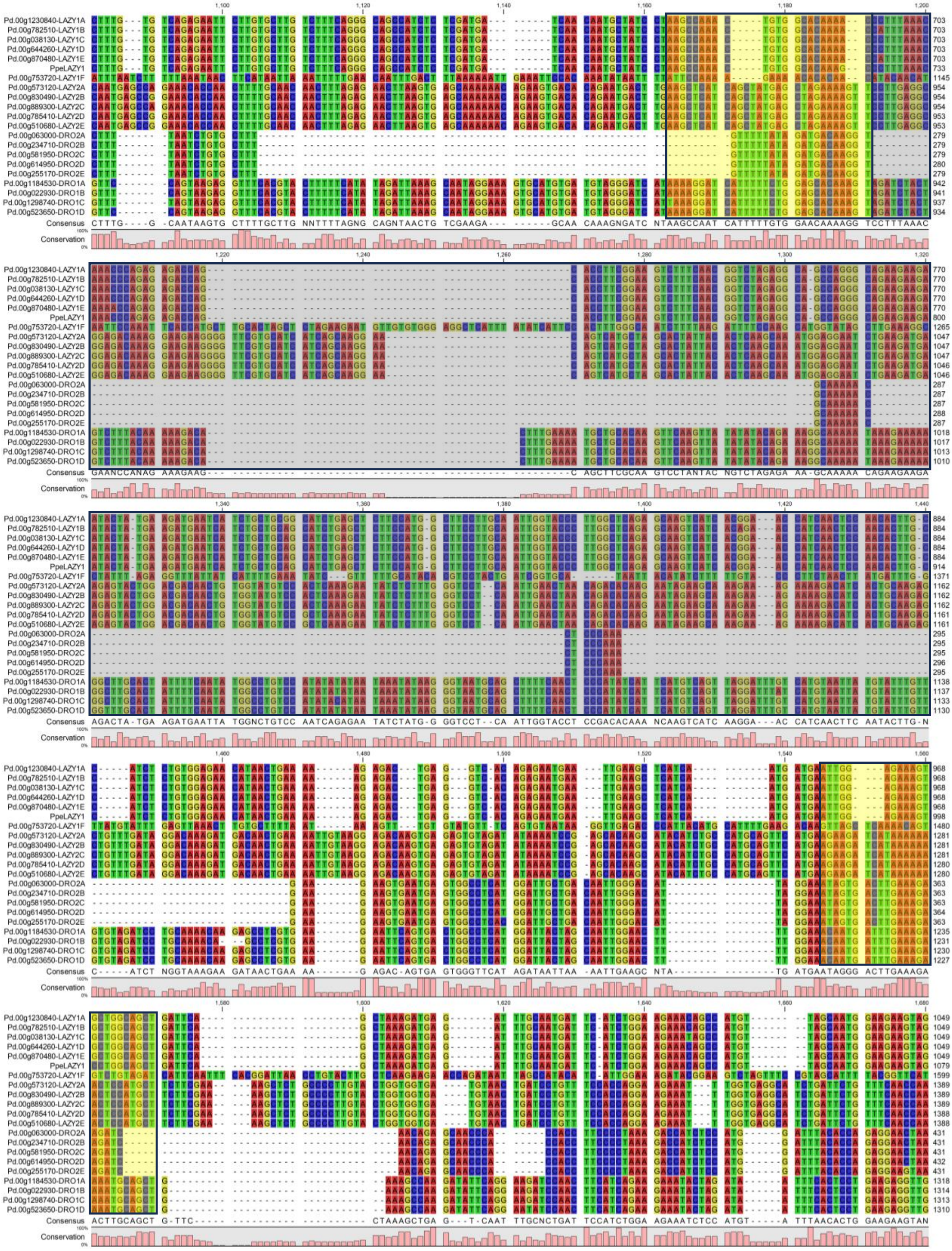

**Figure S3: Plum *LAZY* gene alignment to the insert sequence from *PpeLAZY1*.** Cloning primers highlighted in yellow and the remaining insert sequence in gray.

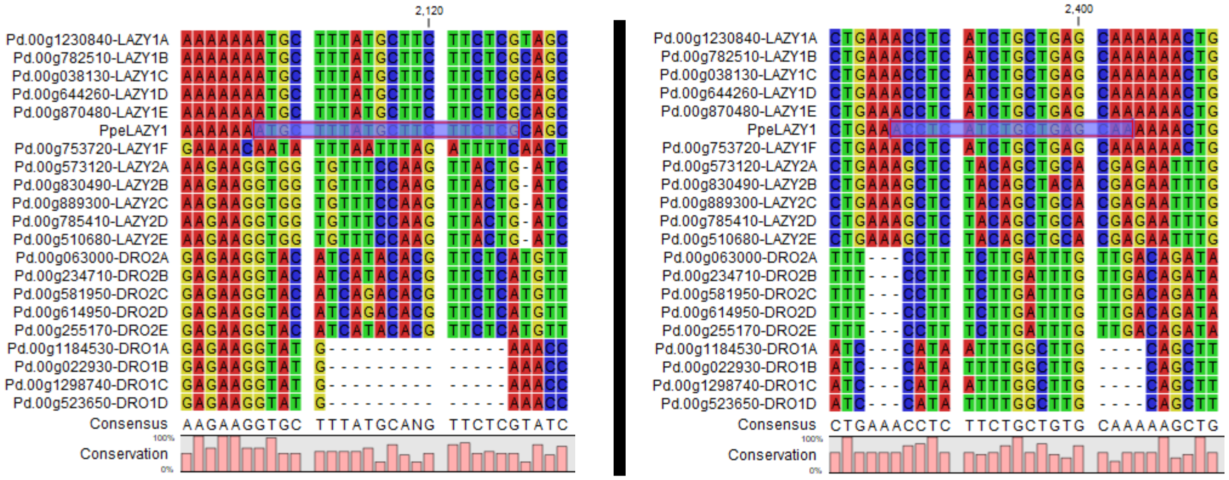


**Figure S4: Sequence alignment for sequence used for gene expression primers.**

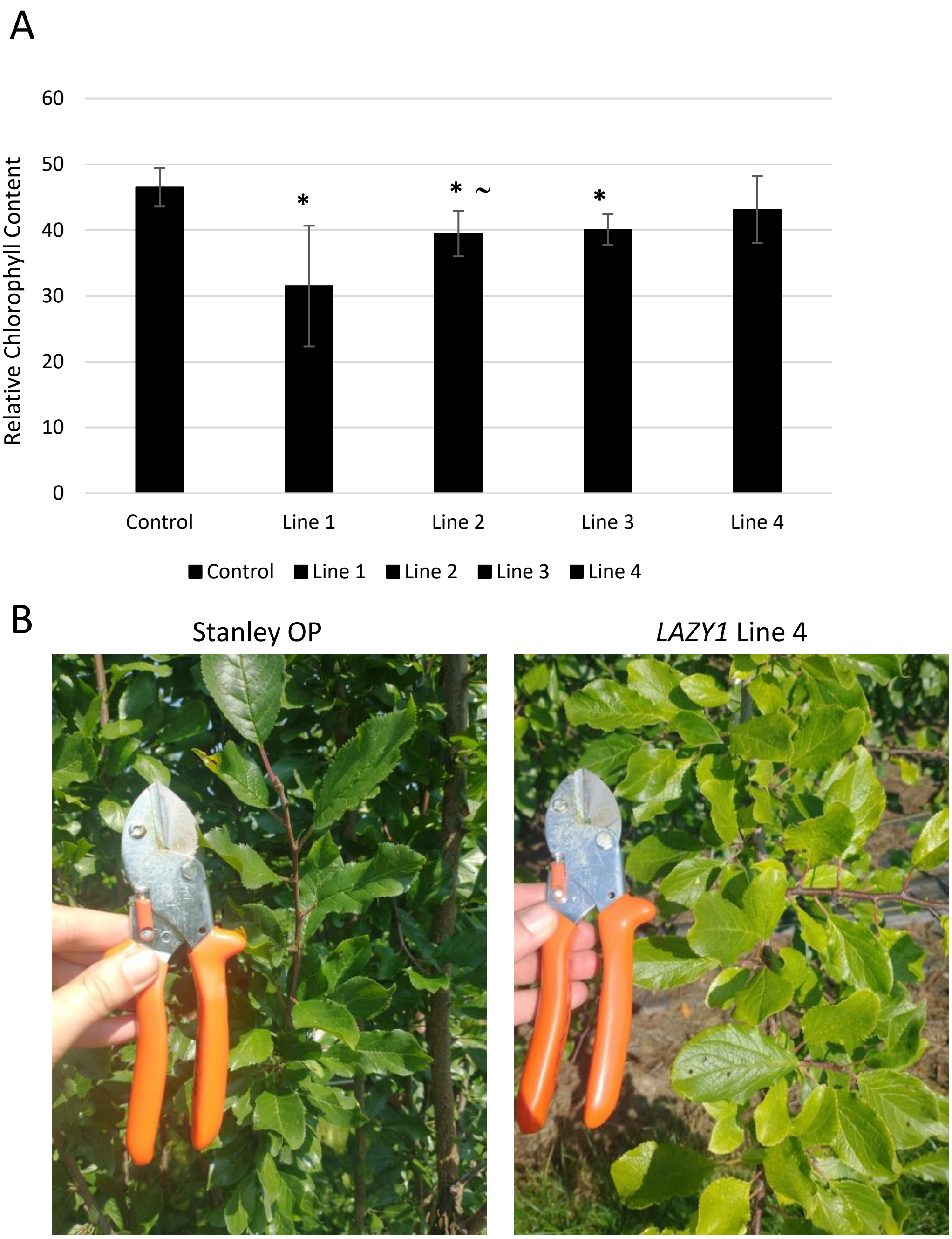


**Figure S5: Additional chlorosis data.** A) 2018 chlorophyll data for WV trees on 8/16/18. ~Line 2 did not have significant reduction in *LAZY1* expression. B) Leaf phenotypes for Stanley OP control and *LAZY1* Line 4 on 7/27/23
